# Agent-Based Modeling of Virtual Tumors Reveals the Critical Influence of Microenvironmental Complexity on Immunotherapy Efficacy

**DOI:** 10.1101/2024.07.03.601920

**Authors:** Yixuan Wang, Daniel R. Bergman, Erica Trujillo, Anthony A. Fernald, Lie Li, Alexander T. Pearson, Randy F. Sweis, Trachette L. Jackson

## Abstract

Since the introduction of the first immune checkpoint inhibitor (ICI), immunotherapy has changed the landscape of molecular therapeutics for cancers. However, ICIs do not work equally well on all cancers and for all patients. There has been a growing interest in using mathematical and computational models to optimize clinical responses. Ordinary differential equations (ODEs) have been widely used for mechanistic modeling in immuno-oncology and immunotherapy because they allow rapid simulations of temporal changes in the cellular and molecular populations involved. Nonetheless, ODEs cannot describe the spatial structure in the tumor microenvironment or quantify the influence of spatially-dependent characteristics of tumor-immune dynamics. For these reasons, agent-based models (ABMs) have gained popularity because they can model more detailed phenotypic and spatial heterogeneity that better reflect the complexity seen in vivo. In the context of anti-PD-1 ICIs, we compare treatment outcomes simulated from an ODE model and an ABM to show the importance of including spatial components in computational models of cancer immunotherapy. We consider tumor cells of high and low antigenicity and two distinct cytotoxic T lymphocyte (CTL) killing mechanisms. The preferred mechanism differs based on the antigenicity of tumor cells. Our ABM reveals varied phenotypic shifts within the tumor and spatial organization of tumor and CTLs, despite similarities in key immune parameters, initial conditions of simulation, and early temporal trajectories of the cell populations.

## 1 Introduction

The versatility of mathematical and computational models has made them an increasingly crucial tool in biomedical research. Models create abstract and simplified representations of real-world phenomena, allowing researchers to gain deeper insights into inherently complex biological processes. These biologically driven and carefully calibrated models extend beyond purely theoretical pursuits. They can shed light on important underlying mechanisms, predict emergent patterns [11], test therapeutic strategies[4], and even inform the design of clinical trials [39, 8]. Immune checkpoint inhibitors (ICIs) are a class of immunotherapeutics that reinvigorate the killing activities of immune cells by blocking the activation of inhibitory immunoreceptors [10]. Immune checkpoint blockade therapy has shown remarkable results for many patients. However, the low overall response rates of ICI monotherapy and the difficulty to enhance patients’ responses with combination therapy in many cancers present an ongoing challenge to clinicians [17, 33]. Adding further complexity to the antitumor immune responses is the fact that cytotoxic T lymphocytes (CTLs) execute their cell-killing function via at least two distinct mechanisms [6, 16]. The first process is mediated by perforin and granzymes. Perforin facilitates the formation of pores in the target cell membrane, which allows granzymes to access the target cell cytoplasm to induce apoptosis [1, 6, 37]. The second process is through the Fas pathway. FasL, a type II transmembrane protein upregulated on CTLs, can engage Fas on the target cell to trigger apoptosis of the target cell [9, 6]. Evidence showed that the perforin/granzyme-mediated process happens faster than the FasL-mediated process [6]. In an in vitro study, perforin-mediated killing completed within thirty minutes, whereas FasL-based killing was detected no sooner than two hours after the tumor cell conjugated with CTL. [9]. Evidence also showed that the switch from fast to slow killing is related to decreasing presence of antigens [19]. Although the connections between distinct CTL killing mechanisms are not fully understood, we find it important to consider the immune system’s varied responses towards tumor cells with different antigenicity and to integrate them into our computational models.

To explain the wide variations of patient responses, quantify the influence of spatial complexity in the tumor microenviroment (TME) , and predict which patients are most likely to respond well to ICIs, we build mathematical and computational models for the ICIs targeting the PD-1/PD-L1 immune checkpoint. Ordinary differential equations (ODEs) and agent-based models are two popular modeling approaches for cancer treatments. An ODE model describes the temporal evolution of populations of cells or molecules through a set of coupled mathematical equations. An ABM simulates how individual entities such as cells and molecules move and interact with each other and with the environment. There have been many ODE-based models in the field of mathematical oncology, including but not limited to works on general tumor-immune dynamics [14], oncolytic virus therapy [13] and anti-PD-1 immune checkpoint inhibitors [24]. Similarly, ABMs have been developed to model the TME and cancer immune response [25]. We previously developed the first ODE model building on the works of [14] and [24], and subsequently the first ABM for anti-PD-1 immune checkpoint blockade therapy with consideration of tumor cells of different antigenicity and the two aforementioned CTL killing mechanisms [36, 2], although the ABM in [2] also includes the anti-FGFR3 small molecule inhibitors. The ABM in this paper is adapted from [2] to focus on the activity of the PD-1/PD-L1 immune checkpoint and the two CTL killing mechanisms like in the ODE model.

The ABM is undoubtedly more complex than the ODE, capturing the phenotypical heterogeneity of the three-dimensional tumor in space, as well as the spatial activities of CTLs in the TME. Our previous work analyzed the ODE model in detail to identify important characteristics of the tumor-immune landscape that have the largest impact on the outcomes of immune checkpoint blockade. Therefore, this paper centers on examining what aspects of the tumor-immune dynamics both the ODE and ABM can describe and what unique insights the ABM can offer due to the integration of the spatial elements. By comparing and contrasting the ABM and the ODE model and using the immune checkpoint inhibitors as an example, we will also discuss the balance between model tractability, model complexity and computational efficiency when building models for cancer immunotherapy.

## 2 Materials and Methods

### Computational models

We compare two mathematical models to describe the tumor-immune dynamics with an active or blocked PD1/PD-L1 immune checkpoint. The first formulation is an ODE model that tracks the temporal changes in the number of tumor cells, CTLs, and concentration of PD-1 and PD-L1. The details of this ODE model are previously published in [36]. The second formulation is a three-dimensional, on-lattice ABM in which tumor cells and immune cells are modeled as autonomous agents interacting with each other and the TME. Like in the ODE model, there are three types of cells in the ABM: high-antigen (HA) tumor cells, low-antigen (LA) tumor cells, and CTLs. Cells in the ABM occupy lattice sites. Tumor cells are immobile while CTLs are mobile. At each time step, tumor cells can proliferate or undergo apoptosis. The proliferation of tumor cells slows down due to contact inhibition [20], because tumor cells are immobile in this ABM. Here we only simulate the virtual tumor until it escapes or metastasizes into nearby blood vessels. Hence, simulations stop when the tumor cells exceed a maximum number allowed or too many tumor cells have reached the boundaries of the TME lattice. The model employs an immune stimulatory factor (ISF), a construct representing the combined effect of factors that each tumor cell secretes into the local neighborhood of the tumor microenvironment. The level of ISF expression depends on the cell’s antigenicity. LA tumor cells secrete a fraction of ISF compared to HA tumor cells.

In the ABM, CTLs are recruited from the lattice boundaries at a constant rate, independent of tumor size. At each time step, a CTL can execute one of the following actions: proliferation, apoptosis or exhaustion, movement, or conjugation. The proliferation rate of CTLs depend on both a base rate and the concentration of ISFs in the surrounding, and is also affected by contact inhibition. CTL exhaustion occurs as a result of extended antigen exposure [18, 5, 22]. CTL apoptosis also arises naturally [22]. Since both dead and exhausted CTLs lose effector functions, the apoptosis and exhaustion of CTLs are combined into a single event in the ABM. The direction of CTL movement is influenced by the concentration gradient of ISF in the TME, i.e., CTLs are more likely to move in the direction of higher ISF. Once CTLs conjugate with a tumor cell, they attempt to destroy it via fast or slow killing. In our previous ABM [2], HA tumor cells are only killed via the fast mechanism and the LA tumor cells are only killed via the slow mechanism. We relax this restriction, adding the probability of fast killing for both HA and LA tumor cells, allowing maximum modeling flexibility and also allowing us to assess the importance of considering the two killing mechanisms in tumor-immune dynamics. The assumption in the baseline parameter set is that CTLs kill HA preferentially via the fast mechanism and kill LA preferentially via the slow mechanism. In both the ABM and the ODE model, an active PD-1/PD-L1 immune checkpoint inhibits the recruitment and antigen-mediated proliferation rates of CTLs in the TME. In both models, we categorize therapeutic outcomes into “elimination”, “dormancy” and “escape”, which corresponds to the three phases of the immunoediting framework [27].

### Description of experiments

For mouse experiments, 6–8 week old female RAG1 KO and C57BL/6J mice were obtained from The Jackson Laboratory. Mice were housed in a specific pathogen-free animal facility at the University of Chicago and used in accordance to the animal experimental guidelines set by the Institute of Animal Care and Use Committee.

The MB49 cell line is a chemical carcinogen-induced urothelial carcinoma cell line derived from a male C57BL/lcrf-a’ mouse. Cells were maintained at 37°C with 5% CO2 in DMEM supplemented with 10% heat-inactivated FCS, penicillin and streptomycin. 1 × 10^6^ MB49 tumor cells were subcutaneously injected into the flank of RAG1 KO (n=27) or C57Bl/6J (n=24). Four types of MB49 cells with different expression levels of the model antigen SIY (SIYRYYGL) were used: Zs green (no SIY), L14 (low SIY), H1 (high SIY) and a mix of L14 and H1 cells with 1:1 ratio. Each type of MB49 cells were injected to five to seven mice of each strain. Mice that died or had tumors with more than 50% ulceration were excluded from the data used for model calibration. Tumors were measured three dimensionally using a digital caliper on Day 7, 10, 12, 14, 17 and 19. Tumor volume was calculated using L x W x H. All mice were sacrificed on Day 20 in accordance with IACUC guidelines for humane endpoints. Tumors were harvested and digested in 10% FBS/RPMI. Single cell suspensions were filtered through a 100 uM cell strainer and stained with antibodies to PD-1, CD69, CD3, CD19, LAG3, Ki67, CD4, CD44, CD45, CD8a, SIY, CD62L, Foxp3, and Live/Dead Viability Dye Zombie NIR. CTLs were analyzed using flow cytometry. The number of CD8 cells is directly measured and the CTL density within the tumor, i.e. the number of CD8 cells per mm^3^ of tumor is calculated. All experimental animal procedures were approved by the University of Chicago Animal Care and Use Committee (IACUC).

### Estimation of model parameters and construction of virtual tumors

To convert tumor volume to number of tumor cells, we assume that 1 mm^3^ of tumor is equivalent to 10^6^ tumor cells [7]. Given this conversion rate and the initial conditions of the experiments and the ABM simulations, each ABM tumor cell represents 50,000 actual tumor cells. The proliferation rate (*α_n_*) and the contact inhibition parameter (*O^prolif^* ) of tumor cells in the ABM were calibrated using the tumor volumes of RAG KO1 mice using the “SMoRe ParS” method developed by [12]. The calibrated parameters are shown in Table 1. In Figure2A, the blue line shows the mean volume of 25 virtual tumors with calibrated *α_n_* and *O^prolif^* . The range of simulated tumor volumes at each time point shows little variation, and the simulated trajectory closely matches the mean tumor volume of RAG KO1 mice, as shown in orange.

**Table 1:** ABM parameters calibrated from experimental data.

| Name | Description | Values (Baseline) |
| --- | --- | --- |
| $\alpha_n$ | Proliferation rate of tumor cells | $2.6 \text{ d}^{-1}$ |
| $O_T^{\text{prolif}}$ | Maximum number of occupied neighbors that still allows tumor cell proliferation (out of 26) | 20 cells |
| $\mu$ | CTL recruitment rate | 2.5 - 25 (8) ABM CTL $\text{d}^{-1}$ |

Fixing the calibrated *α_n_* and *O^prolif^* , we then varied ten other tumor-immune characteristics using Latin Hypercube Sampling in the range given in Table 2 to construct a virtual cohort comprising 12,000 simulated TMEs. Due to the stochastic nature of the ABM and computational limitations, we cannot vary all ABM parameters We chose ten parameters that we believed would have the most impact on therapeutic outcomes based on most sensitive parameters in the ODE model, which describes the same biological process, and our understanding of the spatial components of the ABM. In Figure 2B, the blue line shows the median, interquartile range, and 95% simulated interval of tumor volumes up to Day 19. The orange lines show the mean and standard deviation of tumor volume of C57BL/6J mice on days when measurements were taken. The simulated trajectories lie reasonably close to the experimental data.

**Figure 1:**
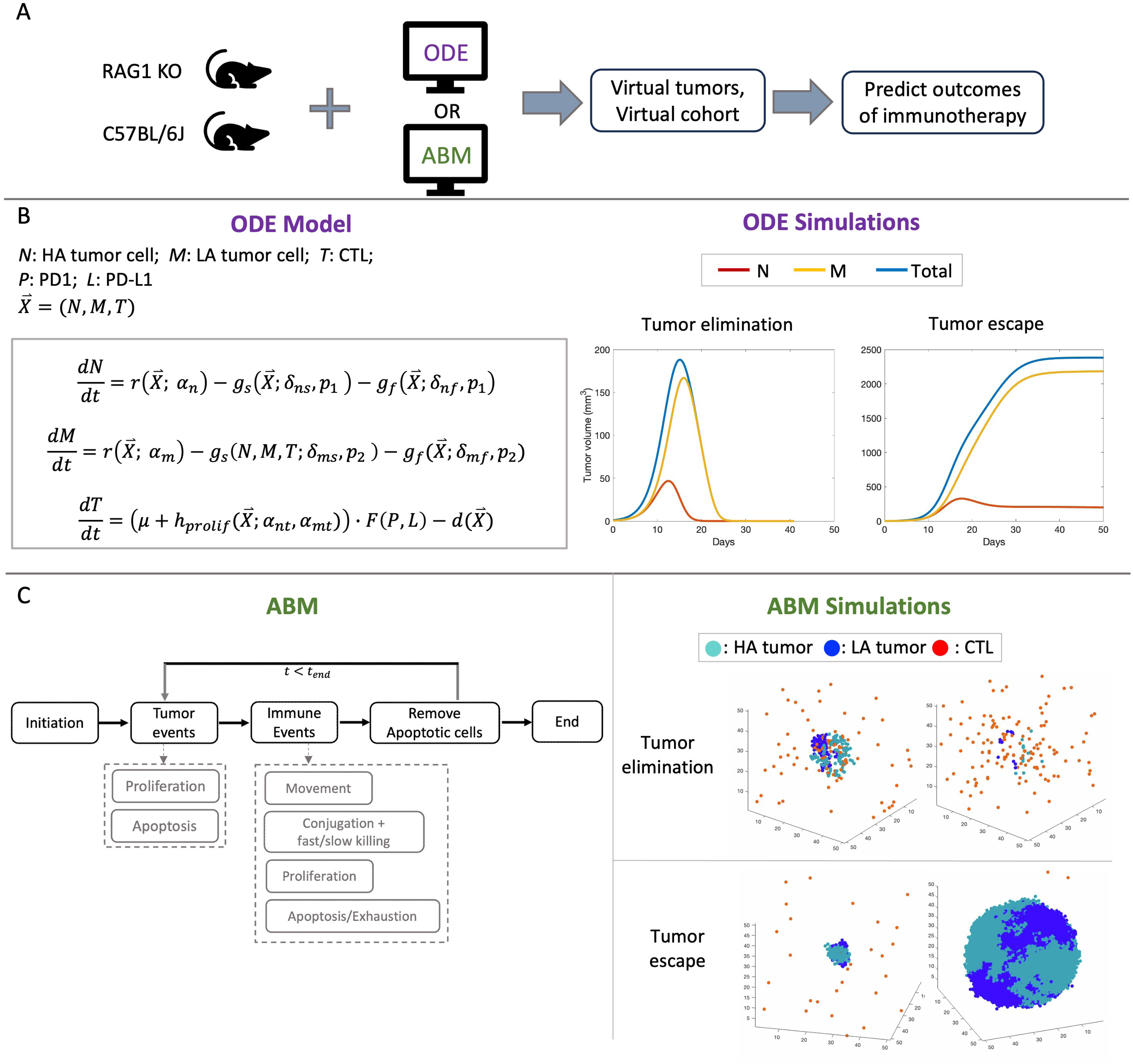
(A) Simulation pipeline: In vivo data of RAG KO1 and C57BL/6J mice calibrate key parameters in the ODE mode and the ABM to simulate virtual tumors and virtual cohort and predict therapeutic outcomes of anti-PD1 immune checkpoint inhibitors (ICIs). (B) Formulation of the ODE model and the temporal trajectories of tumor volumes after immune checkpoint blockade in “elimination” and “escape” cases. HA: high antigen, LA: low antigen. *r*: logistic growth of the tumor; *g_f_* : fast killing of the tumor cells; *g_s_*: slow killing of the tumor cells; *μ*: constant recruitment of CTLs per day; *h_prolif_* : antigen-stimulated proliferation of CTLs; *F* : immune suppression by the PD1-PD-L1 complex; *d*: death/exhaustion of CTLs. See [36] for exact formulation of the ODE model. See S.Table 2 for descriptions of key ODE parameters shown in this figure. (C) Simplified flowchart of the ABM and simulations of tumor elimination and tumor escape after immune checkpoint blockade . At each time step, each tumor cell can either proliferate or undergo apoptosis. Each CTL undergoes one of the four events: movement, conjugation with tumor cells to attach via fast or slow killing, proliferation, apoptosis or exhaustion.

**Figure 2:**
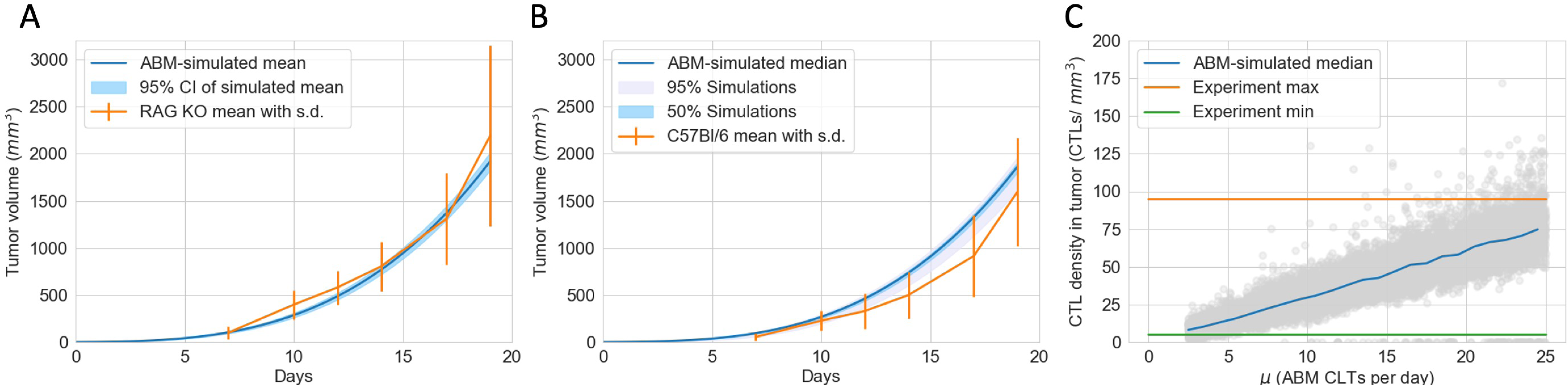
The ABM is calibrated to closely reflect experimental data of mice with an active PD-1/PD-L1 immune checkpoint. (A) Tumor volumes of immunocompromised RAG KO1 mice (in orange) and virtual tumors in the absence of an immune system (in blue) from Day 0 to Day 19. Error bar: standard deviation of RAG KO1 tumor volumes. Shaded blue region: 95% confidence interval of the simulated mean. (B) Tumor volumes of immunocompetent C57Bl/6J mice (in orange) and a virtual cohort comprising 12000 simulated tumors with immune responses (in blue) from Day 0 to Day 19. Error bar: standard deviation of CB57Bl/6J tumor volumes. Shaded grey region: 2.5 to 97.5 percentile of simulated tumor volumes on each day. Shaded blue region: 25 to 75 percentile of simulated tumor volumes on each day. (C) Grey circles: simulated endpoint CTL densities in virtual tumors at different CTL recruitment rates (*μ*). Blue line: median CTL density of virtual tumors with *μ* values in each integer bin. Orange and blue lines: maximum and minimum CTL densities in experimental data of CB57Bl/6J mice on Day 19.

**Table 2:** Parameters varied in the ABM.

| Name | Description | Values (Baseline) | Source |
| --- | --- | --- | --- |
| $\mu$ | CTL recruitment rate | 2.5 - 25 (8) ABM CTL $\text{d}^{-1}$ | Calibrated |
| $ha_0$ | Initial proportion of HA cells | 0.05 - 0.95 (0.5) $\text{d}^{-1}$ | Estimated [2] |
| $\alpha_{nt}$ | Max ISF stimulated CTL proliferation rate | 0.04 - 1.00 (0.15) $\text{d}^{-1}$ | Estimated [14, 26, 24] |
| $\delta_{\text{fast}}$ | Fast kill rate | 12 - 120(48) $\text{d}^{-1}$ | Estimated [2] |
| $p_1$ | Probability of fast killing for HA | 0 - 1(0.92) | Assumed |
| $p_2$ | Probability of fast killing for LA | 0 - 1 (0.33) | Assumed |
| $m$ | CTL movement rate | 1 - 8 (2) $\mu\text{m min}^{-1}$ | Estimated [2] |
| $\beta$ | CTL Conjugation rate with tumor cells | 0.5 - 4 (1.2) $\text{h}^{-1}$ | [15, 2] |
| $a_{\text{reach}}$ | Immune stimulatory factor reach | 60 - 200 (100) $\mu\text{m}$ | Estimated [2] |
| $r_{\text{ISF}}$ | ISF expression by LA tumor cells compared to HA tumor cells | 0.1 - 0.9 (0.5) | Assumed [2] |

**Table 3:** Initial conditions.

| Name | Description | Value |
| --- | --- | --- |
| $N_0$ | Total Tumor cells | 20 ABM cells |
| $T_0$ | CTLs | 0 ABM cells |
| $ha_0$ | Ratio of HA tumor cells to total tumor cells | 0.05 - 0.95 |

Based on the calculated density of CTLs within the tumor in C57BL/6J mice at the endpoint of Day 19, we estimated each ABM CTL represents 2175 actual CTL cells. This scale was calculated, and the range of the CTL recruitment rate in the ABM was chosen so that the range of simulated CTL densities on Day 19 in the virtual cohort matches the range observed experimentally, as illustrated in Figure 2C. The green and orange lines show the observed minimum and maximum CTL density in C57BL/6J mice on Day 19. The grey dots show the endpoint CTL density of each virtual mouse, and the blue line shows the median CTL density for each integer interval of CTL recruitment rate (e.g., 2-3, 19-20, etc.).

## 3 Results

### 3.1 Immunotherapy Efficacy Widely Varies in Virtual Cohort with Indistinguishable Pretreatment Tumor Growth Patterns

To explore the best-case scenarios of checkpoint blockade therapy in the same virtual cohort as in Figure 2B for which pre-treatment growth patterns are similar, we simulated completely blocking the PD-1 PD-L1 immune checkpoint in both the ODE model and the ABM. Figures 3A and B show the median, 95% and 50% simulated interval of tumor volume, with A corresponding to the ODE simulations and B corresponding to the ABM simulations. Both ODE and ABM are able to capture a wide range of treatment outcomes after immune checkpoint blockade, as shown by similar 95% simulated interval (shaded grey) and 50% simulated interval (shaded light blue). Tumor status after treatment ranges from elimination to escape by Day 19. This result contrasts with the tight interval of simulated tumor growth in Figure2B, for the same virtual cohort of mice with an active immune checkpoint. This implies that tumors that grow similarly pre-treatment can have drastically different therapeutic outcomes after immune checkpoint blockade therapy.

**Figure 3:**
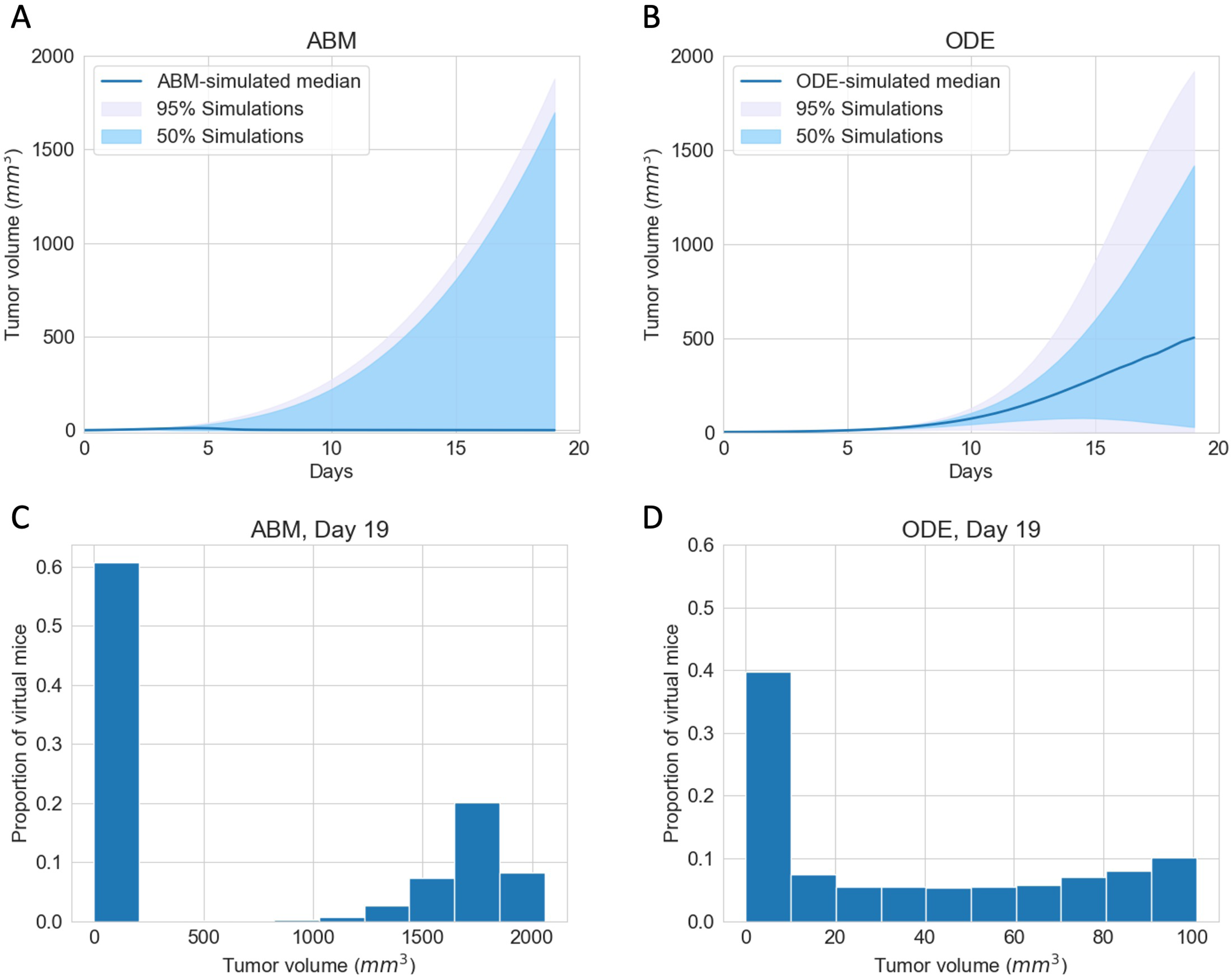
ABM and ODE virtual cohort simulations show a wide range of efficacies for immune checkpoint blockade therapy. (A) ABM simulations of virtual cohort response to ICI. (B) ODE simulations of virtual cohort response to ICI. Blue line: simulated median. Shaded grey region: 2.5 to 97.5 percentile of simulated tumor volumes on each day. Shaded blue region: 25 to 75 percentile of simulated tumor volumes on each day. (C) Histogram of ABM-simulated tumor volumes on Day 19. (D) Histogram of ODE-simulated tumor volumes on Day 19.

We notice stark differences in the median trajectories of the tumor volume in (A) and (B). In the ODE model, the median tumor volume achieves a moderate size, which we characterize as dormancy, by Day 19; whereas in the ABM, the median tumor volume is small enough to be described as eliminated by Day 19. A closer look at tumor volumes on Day 19 reveals that most tumors resolve into an elimination or escape steady state outcome much faster in the ABM than in the ODE. Figures 3C and D show the distribution of tumor volume on Day 19. Tumors in ODE simulations range from 0 to 2000 mm^3^, whereas tumors in the ABM simulations are either close to mm^3^ or above 1000mm^3^, with no in-between cases. The lack of intermediate tumor sizes on Day 19 in the ABM suggests that a typical virtual tumor either gets eliminated or escapes by Day 19.

### 3.2 Initial phenotypic composition dictates composition and volume of tumor after checkpoint blockade therapy

In virtual clones with identical tumor and immune characteristics, ODE and ABM simulations show that different initial percentages of LA tumor cells result in different outcomes of checkpoint blockade therapy. The baseline assumption of our models is that LA tumor cells have a survival advantage over HA tumor cells. In the ODE model and the ABM, HA tumor cells are more likely to be killed via the fast mechanism, and LA tumor cells are more likely to be killed via the slow mechanism. Moreover, in the ABM, CTLs are more likely to move towards HA tumor cells than LA tumor cells, increasing the likelihood of CTL conjugation with HA tumor cells and HA cell clearance.

For the chosen set of parameters, Figure 4A shows that, using the ODE model, if the initial tumor is 50% or 80% LA tumor cells, the tumor grows to the maximum possible volume. Using the ABM model, Figures 4B and C show that the tumor escapes, but there are slight variations in the long-term steady-state tumor volume. This variation in final tumor volume in panels B and C is likely due to one of the stopping criteria of our ABM. The ABM stops and marks the tumor as “escape” once sufficient ABM tumor cells reach the boundary of the TME, and this might happen at different times for each of these tumors that escaped. Therefore, the tumors in panels B and C have no qualitative differences in terms of tumor volume. With 20% LA tumor cells initially in the ODE model, the tumor gets eliminated by Day 28. In the ABM, an initial LA ratio of 20% makes tumor elimination possible but not guaranteed. In this case, the outcome ranges from tumor elimination to tumor escape across simulations, with a wide range of possible steady-state tumor sizes in between.

**Figure 4:**
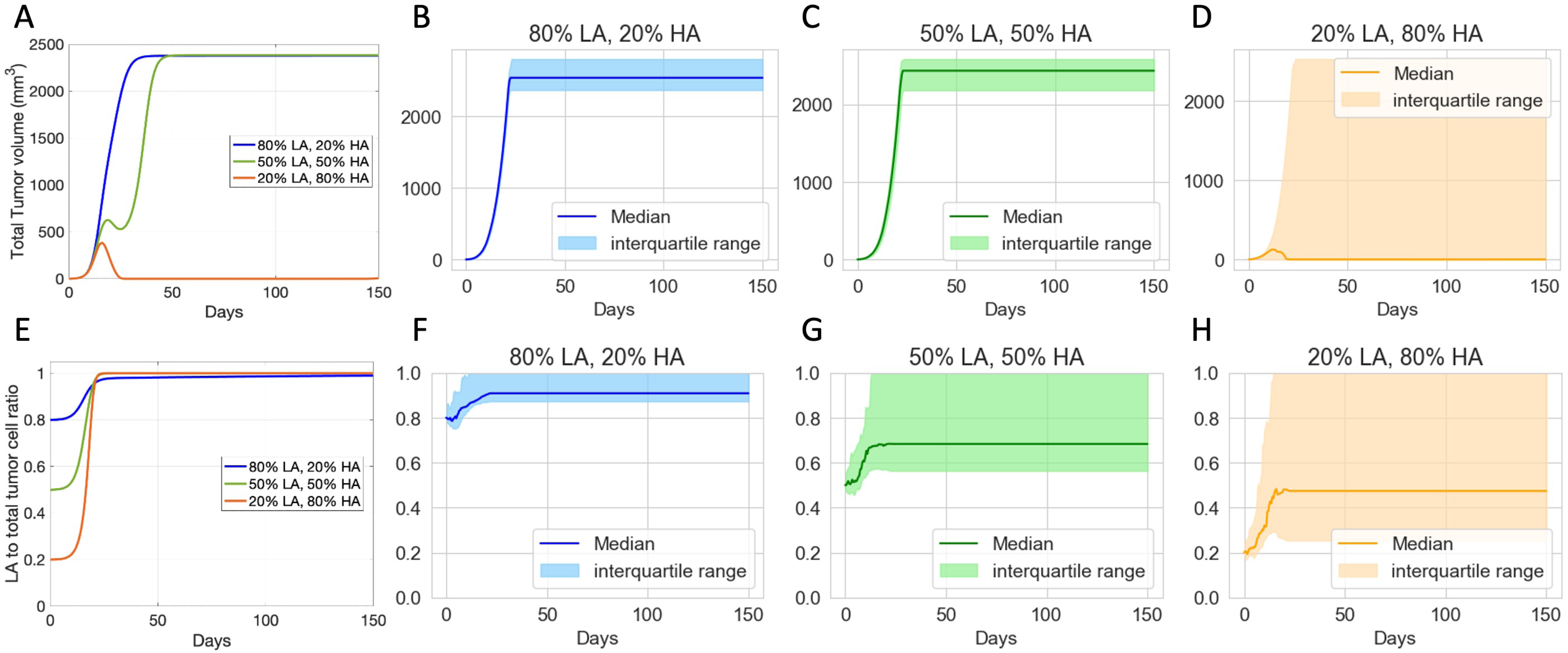
Virtual clones with identical tumor-immune characteristics but different initial tumor composition show varied long-term response to checkpoint blockade therapy. (A) Volumes of ODE-simulated virtual tumors. (B-D) Volumes of ABM-simulated virtual tumors. (E) Ratio of low-antigent (LA) tumor cells to total tumor cells in ODE-simulated virtual tumors. (F-H) Ratio of LA tumor cell to total tumor cell in ABM-simulated virtual tumors. Colors show different compositions of the initial tumor. Blue: 80% LA tumor cells, 20% HA tumor cell. Green: 50% LA, 50% LA. Orange: 20% LA, 80% HA. ODE parameters for virtual clones in (A) and (E): *α_nt_* = 0.32, *μ* = 2×10^4^. ABM parameters for virtual clones in (B-D) and (F-H): *α_nt_* = 0.32, *μ* = 15. Other parameters are set at baseline.

In terms of tumor composition, across all ODE and ABM simulations with different initial tumor compositions, the final tumor is always more LA-dominant than the initial tumor, even in the cases where the tumor shrinks. This result reflects the survival advantages that LA cells have in our models. It also suggests that checkpoint blockade therapy can reduce the tumor size but increase the proportion of LA cells in the resulting tumor, which can impact the results of other subsequent immunotherapies. While both ODE and ABM simulations show a trend toward increased LA ratio, there are notable differences in the final tumor compositions. In the ODE simulations, the final tumor is consistently 100% LA tumor cells. However, in the ABM, the final tumor can exhibit a wider range of LA ratios, with the range expanding as the initial ratio of LA tumor cells decreases.

### 3.3 Therapeutic outcomes are correlated with key immune parameters

We explore what tumor or immune characteristics are most correlated with therapeutic outcomes of checkpoint blockade in the ODE model and the ABM model. In particular, we are interested in knowing whether there are similar conclusions for parameters common to both models and how spatial parameters unique to the ABM relate to the outcomes. In our extensive analysis of the ODE model [36], we determined that the CTL recruitment rate (*μ*) is the most important immune parameter for achieving tumor reduction and elimination. The ABM indeed corroborates this result. Figure 5A shows the distribution, median, and interquartile range of the CTL recruitment rate in the ODE model in the virtual cohort associated with each therapeutic outcome. Figure 5B shows a similar graph for *μ* in the ABM. The median and interquartile range of *μ* associated with tumors that eventually get eliminated are higher than those of escape cases. The ABM shows more prominent separation of the distribution of *μ* between elimination and escape cases, as the 75 percentile in the escape cases is lower than the 25 percentile in the elimination cases. This suggests the CTL recruitment rate might be even more predictive of treatment outcomes in the ABM than in the ODE model.

**Figure 5:**
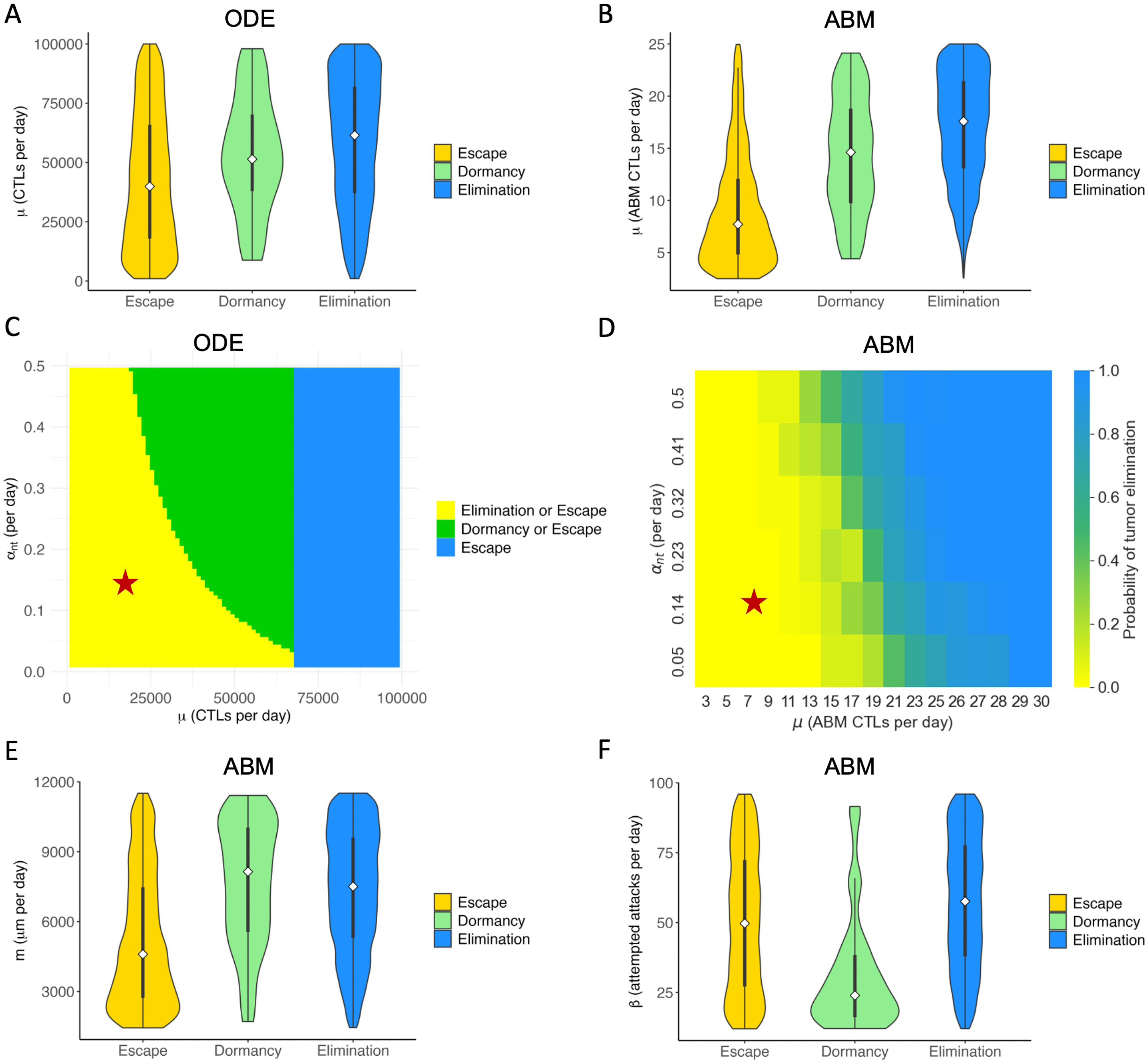
Key immune parameters in ABM and ODE models are correlated with outcomes of immune checkpoint blockade. Colors in all sub-panels represent long-term (*t* ≥ 150 days) outcomes of checkpoint blockade therapy. Elimination (blue): tumor size *<* 0.1mm^3^; Dormancy (green): *<* tumor size *<* 500mm^3^; Escape (yellow): tumor size *>* 500mm^3^. (A) ODE: Violin plot of the distribution of CTL recruitment rate (*μ*) in the virtual cohort associated with each outcome, with the shape showing probability density, the white circle showing the median, and the black lines showing the interquartile range. (B) ABM: Violin plot of the distribution of *μ* in the virtual cohort associated with each outcome. (C) ODE: Two-parameter bifurcation diagram of *μ* and max antigen-stimulated CTL proliferation rate (*α_mt_*) on steady state tumor size. Red star: baseline parameters. (D) ABM: Probability of tumor elimination at each *μ*-*α_nt_* combination. Colormap shows the probability of tumor elimination ranging from 0 (yellow) to 1 (blue). (E) ABM: CTL movement rate (*m*). (F) ABM: CTL conjugation rate (*β*).

In our analysis of the ODE model, we examined the combined effect of varying both CTL recruitment rate and maximum antigen-mediated CTL proliferation rate. The parameter with an equivalent effect in the ABM is the maximum ISF-stimulated proliferation rate of CTLs (*α_nt_*). Figure 5C shows the two-parameter bifurcation diagram of *μ* and *α_nt_* in the ABM. An analogous approach was used in the ABM to capture the effect of the ODE bifurcation analysis. Figure 5D depicts at the probability of tumor elimination in the virtual cohort of mice as *μ* and *α_nt_* vary. Blue represents a 100% chance of elimination, and yellow represents a 0% chance of tumor elimination. Due to the stochastic nature of the ABM, different runs of the same set of parameters yield different tumor outcomes, resulting in the gradient of colors between blue and yellow in panel D. Green regions Figure 5D thus represents a non-zero probability of tumor elimination. Both C and D show that our baseline case lies in the region of parameter space where the tumor escapes to carrying capacity with certainty. For virtual tumors with a low *α_nt_*value, only increasing *α_nt_*is ineffective for reducing tumor volume in the long term. The most efficient way to reduce tumor volume at equilibrium is by increasing *α_nt_* and *μ* simultaneously, landing in the green region where the tumor can be eliminated. Increasing *μ* sufficiently can eliminate the tumor with certainty.

Figure 5E and F show the distribution of spatial parameters of tumors associated with each outcome. Figure 5E shows that escape cases generally have a much lower T cell movement rate (*m*) than elimination and dormancy cases. A low T cell movement rate, as depicted in Figure 5 E, has significant consequences. It hampers the ability of CTLs to reach tumor cells fast enough to prevent the tumor from spreading to the lattice’s edge and escaping. Moreover, in the ABM cohort, only a small number of tumors have not been eliminated or escaped by Day 150, therefore being marked as dormant. Figure 5F shows that most dormancy cases have a low conjugation rate (*β*) between tumor and CTLs. The low conjugation rate is a significant factor in tumor dormancy. It leads to reduced interaction, prolonging the time to reach a stable term equilibrium, as shown in Figure 5F. We observe a similar pattern with the LA-ISF factor, the ratio of ISF secreted by LA tumor cells compared to HA tumor cells, as seen in Figure S1. As CTLs are more likely to move towards regions with high ISF, a low LA-ISF factor makes it harder for CTLs to locate and conjugate with LA cells. Therefore, low *β* has a similar effect as a low *m*: both cause the tumor to stay dormant for longer.

### 3.4 ABM reveals spatial and phenotypic heterogeneity despite similar temporal tumor and immune growth patterns

To explore what additional insights we gain using an ABM that includes spatial features, we examine tumors that are expected to grow similarly after checkpoint blockade in the ODE model but end up having different or even completely opposite therapeutic outcomes in the ABM. In particular, we analyze 603 virtual tumors with the same number of tumor cells and CTLs initially and similar CTL temporal trajectories up to Day 7. Eventually, 251 of these tumors are eliminated, and 352 escape long-term. Based on their future tumor status, we categorize these 603 tumors as “to be eliminated” and ‘ ‘to escape,” or “elimination” and “escape,” in short. Figure 6A shows that despite the same initial condition, the number of tumor cells in the “elimination” and “escape” groups diverge quickly between Day 2 and Day 3. However, the total number of CTLs in the TME in both groups remains close up to Day 7, as illustrated in Figure 6B. Analysis of the ODE model in [36] identified CTL recruitment rate as the most critical parameter for predicting tumor outcomes after checkpoint blockade, followed by maximum antigen-stimulated CTL proliferation rate and fast-kill rate. Figures 6C,D,E show similar distributions and close medians of these key immune parameters in “elimination” and “escape” groups. In Figure 4, we saw that initial tumor compositions can greatly influence the outcomes of checkpoint blockade therapy. Nonetheless, there is no clear patterns of LA-dominance or HA-dominance in the initial tumors in either group, as illustrated by comparable distributions and close medians of initial ratio of HA tumor cells (*ha*_0_) in Figure 6F. The probability of fast killing (*p*_1_*, p*_2_) were not identified as sensitive parameters for tumor volumes in the ODE model [36]. Nonetheless, here in the ABM, they show some correlation with the therapeutic outcomes as seen in Figure S2. Tumors with low *p*_1_ are more likely to escape. Other relationships between the probability of tumor elimination and the values of *p*_1_ and *p*_2_ are weaker. Given similar initial conditions, critical immune parameters, and similar temporal trajectories of the number of CTLs, the ODE model is unable to describe or explain the disparate treatment outcomes. Therefore, we focus on parameters and features that are unique to the ABM next.

**Figure 6:**
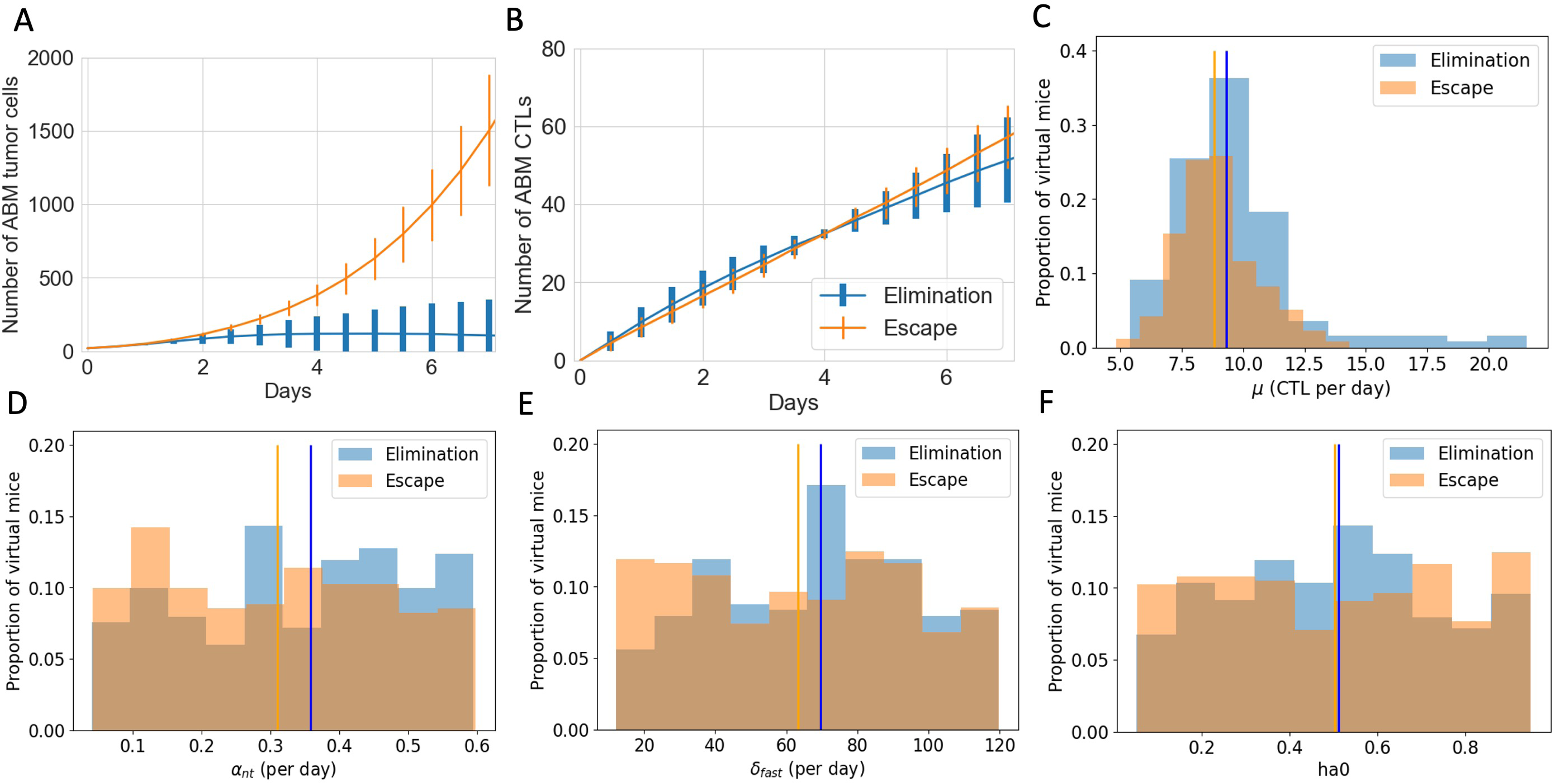
Virtual tumors with similar number of CTLs in the TME up to Day 7 show diverging tumor control outcomes after checkpoint blockade. Blue: tumor elimination; orange: tumor escape. (A) Number of ABM tumor cells up to Day 7. Lines show the mean and error bars show the standard deviation of tumor volume in each group. (B) Number of ABM CTLs up to Day 7. (C) Histogram of CTL recruitment rates (*μ*) in “elimination” and “escape” groups respectively, with the y-value normalized by the total number of virtual tumors in that group. Vertical lines show the median *μ* of virtual tumors in each group. (D) Max ISF-stimulated CTL proliferation rate (*α_nt_*) (E) Rate of fast killing by CTLs (F) Initial ratio of HA tumor cells to total tumor cells.

Figures 6B shows the temporal evolution of the total number of CTLs in the TME. However, it does not tell us where these CTLs are and how they interact with the tumor cells. Figures 7 complements information about CTLs in the ABM. Figure 7A shows that despite the “escape” group and “elimination” group having similar numbers of CTLs, there are clearly more tumor cells cleared by CTLs per day in the “elimination” group. Figure 7B shows that on average, at each time point, CTLs in the “elimination” group are closer to the tumor center than those in the “escape” group in the first four days, allowing CTLs to be closer to tumor cells and have a higher chance of clearing tumor cells early on. Another spatial feature is the volume of the tumor convex hull, which is defined as the smallest convex shape that includes all the tumor cells. Figure 7C shows a smaller mean tumor convex hull in the “elimination” group, meaning that the tumor is more compact. The combined effect of more CTLs closer to the tumor center and a smaller convex hull is that the CTL density within the tumor is higher in the “elimination” group starting on Day 2, making conjugations and tumor cell clearance more likely. In addition to examining CTLs in relation to the tumor center, we also consider the spatial distribution of CTLs with respect to each tumor cell. Figure 7D and E show the temporal evolution of the distributions of CTLs relative to tumor cells in the ABM. The top-down direction shows the evolution of time. The x-axis shows the distance from a tumor cell. The color map represents the mean number of CTLs at a certain distance from a tumor cell at each time point from Day 0 to Day 7, averaged across all virtual tumors in each group. In the first four days, there are more CTLs close to tumor cells (e.g., distance *<* 160*μm*) in the “elimination” group. From Day 4 onwards, although the number of CTLs near each tumor cell increases in the “escape” group, it was too late. The tumors still escape eventually.

**Figure 7:**
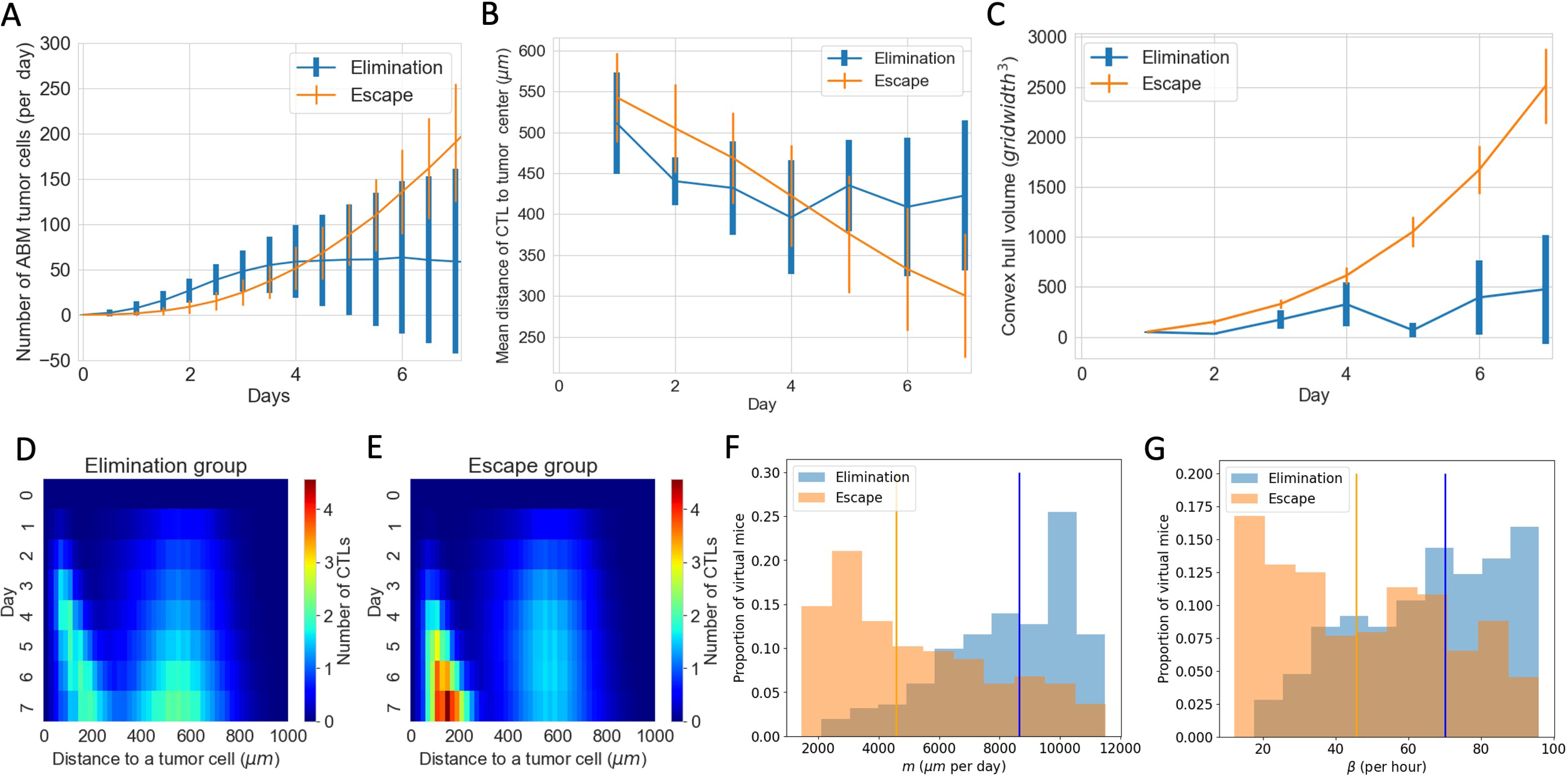
Spatial features explain diverging therapeutic outcomes after immune checkpoint blockade despite similar total number of CTLs in the TME. (A) Number of tumor cells cleared by CTLs per day in “elimination” and “escape” groups. Blue: elimination; orange: escape. The line shows the mean and error bars show standard deviation in each group. (B) Mean distance of CTLs to the tumor center. (C) Volume of the tumor convex hull (D,E) Y-axis shows time. X-axis shows the distance of a CTL to a tumor cell. The color represents the mean number of CTLs at a certain distance from a tumor cell at that time point. D corresponds to the “elimination” group and E corresponds to the “escape” group. (F) Normalized histogram of CTL movement rate (*m*) of virtual tumors in “elimination” and “escape” groups respectively. Vertical lines: median *m* in each group. (G) CTL conjugation rate (*β*).

Parameters existing in both the ABM and the ODE model show no clear pattern in distribution in “elimination” and “escape” groups in Figure 6. However, spatial parameters in the ABM that are not captured by the ODE model show distinct distribution patterns corresponding to each outcome group. This helps explain the observations about number of tumor cells cleared per day and spatial distributions of CTLs in the “elimination” and “escape” groups in Figure 7 A, B, D and E. Movement rate (*m*) and conjugation rate (*β*) skew to the right in the “elimination” group and to the left in the “escape” group in Figure7F and G. Almost all virtual tumors with extremely low *m* or *β* values escape. On the contrary, a virtual tumor with high *m* or *β* values are much more likely to get eliminated than to escape. CTLs with high *m* values have higher motility, and are thus able to collocate to tumor cells faster. CTLs with high *β* values are more likely to conjugate with tumor cells at each time step. Therefore, given a similar number of CTLs, virtual tumors with higher CTL movement and conjugation rates are more likely to have favorable outcomes after checkpoint blockade therapy in the long term.

## 4 Discussion

Here, we present a comparison of an ODE model with an ABM for the same cancer immunotherapy: ICI for the PD-1/PD-L1 immune checkpoint. We simulate tumor progression and the response to immune checkpoint blockade therapy in a virtual cohort using a three-dimensional, on-lattice ABM calibrated using in vivo data from bladder cancer studies in mice. Our models reveal which tumor and immune characteristics affect the outcomes of checkpoint blockade therapy the most. While our previous work [36] analyzed the ODE models thoroughly, this paper focuses on the capabilities of the ABM. In this way, we explore what biological insights both models can provide and what additional insights the ABM offers about the spatial complexity of the TME and its impact on therapeutic outcomes. Despite the enhanced modeling capabilities, the use of ABMs also presents challenges. Therefore, we will also discuss the pros and cons of the ODE model and the ABM for modeling tumor-immune dynamics.

The ODE model and ABM predict a wide range of therapeutic responses to immune checkpoint blockade therapy in a virtual cohort with similar tumor growth pre-treatment. Both models also identify crucial immune parameters linked to the range of outcomes. Our analysis of both models underscores the pivotal role of CTL recruitment rate (*μ*) and maximum rate of antigen-mediated CTL proliferation (*α_nt_*) in tumor reduction or elimination. Since adoptive T cell therapy can increase *μ* and therapeutic cytokines like interleukin-2 (IL-2) can increase *α_nt_*, our results in Figure 5 A-D have implications on the effectiveness of combination therapy strategies. Our simulations suggest that combination therapy of anti-PD1 and adoptive T cell transfer is effective in reducing tumor size drastically or eliminating tumors if CTL recruitment rate can be enhanced to sufficiently high levels. Various combination therapy strategies involving anti-PD1 and tumor-infiltrating lymphocytes (TIL) or chimeric antigen receptor (CAR) T cells have shown synergistic effects in both preclinical studies and clinical trials [28, 29]. Lifileucel, a TIL therapy was recently approved for patients who received prior treatment with anti–PD1/PD-L1 antibodies [29]. Our simulations suggest another possible way to achieve a drastic reduction of tumor volume or even tumor elimination with a smaller amount of drug: combining ICIs with both adoptive T cell transfer and cytokine-directed therapy. In this way, a patient’s parameters can move from a baseline outcome of ICI monotherapy, where the tumor escapes with certainty, to a region in parameter space where tumor elimination is possible. In fact, IL-2 treatments are often administered with other forms of immunotherapy, such as Lifileucel. Furthermore, IL-2 therapy in combination with anti-PD1/PD-L1 was shown to be feasible and tolerable, although clinical trials to show effectiveness of this therapy are still underway [30, 32].

Both models also show the importance of considering tumor antigenicity and multiple immune-cell kill mechanisms preferentially associated with HA or LA tumor cells. Our baseline assumptions was that CTLs preferentially kill HA cells via the fast mechanism and LA cells via the slow mechanism. Effectively, we assumed that LA cells are the harder-to-treat phenotype regarding antigenicity. Using virtual clones with different initial LA to total tumor cell ratio, we showed that the less LA-dominant the initial tumor is, the better the outcomes after immune checkpoint blockade. Moreover, the final tumor was always more LA-dominant than the initial tumor. These are both consequences of CTLs killing HA tumor cells faster than LA tumor cells. Higher numbers of LA cells in the resulting tumor suggests that if ICI does not eliminate the tumor, it might become a “colder” tumor, thereby affecting the responses to subsequent treatments. The shift to LA-dominance aligns with well-documented observations of immune selection for lowly antigenic tumors [38]. In the ABM, the antigenicity of the tumor cells not only determines how fast CTLs kill tumor cells once conjugation has occurred, it also greatly impacts the movement of CTLs before conjugation. CTLs gravitate towards regions with high ISF, and HA tumor cells secret higher ISF in our model. This key difference between HA and LA tumor cells underlies the impact of the CTL movement rate, conjugation rate and LA-ISF factors on treatment outcomes.

The ABM enhances our understanding of the TME by incorporating spatial characteristics that ODEs cannot capture. This allows for more nuanced insights, revealing complexities that might be overlooked when immune parameters, initial tumor composition, and the temporal evolution of cellular populations appear similar. In both models, we observed the importance of CTL recruitment rate (*μ*) and max antigen-stimulated CTL proliferation rate (*α_nt_*) to tumor elimination after immune checkpoint blockade. This might seem intuitive as higher *μ* and *α_nt_* results in more active CTLs in the TME, and thus, they eliminate more tumor cells. However, we chose 603 virtual tumors from ABM simulations to show that, when considering intratumoral spatial heterogeneity, tumors with similar *μ* and *α_nt_*values and similar temporal trajectories of CTLs in the TME can experience drastically different fates after checkpoint blockade therapy (Figure 6, 7). In the ODE model formulation, CTLs indiscriminately target all HA or all LA. By contrast, in the ABM, immune attacks are contingent on CTLs moving toward tumor cells and successfully conjugating with them. Therefore, the movement rate of CTLs (*m*) and the conjugation rate of CTLs with tumor cells (*β*) prove to be crucial in determining how fast CTLs colocalize to, attack and clear tumor cells. Virtual tumors with high *m* and *β* are more likely to get eliminated after checkpoint blockade. Translational data are emerging on the critical nature of spatial relationships in the immune tumor microenvironment. A multitude of factors such as gradients of chemokines and physical features of the microenvironment have been shown to affect T cell movement [35]. In a melanoma mouse study, adoptive T cell therapy successfully controled tumor growth in some cases but failed in others. The T cell infiltration and motility were higher in responders relative to non-responders, as evidenced by increased speed and distance traveled of T cells [31]. An in vitro study of melanoma showed that varied ICI responses were not merely due to differences in tumor structure or proportion of cell types. Physical proximity and contact frequency between CTLs and tumor cells also significantly differed between responders and nonresponders of ICIs [21]. Among many ongoing efforts to develop therapeutics to enhance the T cell motility and infiltration, tebentafusp, a bispecific protein consisting of an affinity-enhanced T cell receptor fused to an anti-CD3 effector that can redirect T cells to target glycoprotein 100–positive cells [23], was FDA-approved in 2022.

With proper formulation, both ABMs and ODE models can accurately reflect biological processes. Still, they have a few fundamental differences, which lead to their respective pros and cons from the modeling perspective. ODEs model the population-level temporal dynamics of each type of cell or drug molecule, whereas ABMs model each cell as an autonomous agent. At a given time point, all cells or molecules of a single type in the ODE undergo the same changes uniformly, whereas agents in the ABM experience different events based on their location in space or what other agents surround them. Thus, the ABM is more flexible in modeling intratumoral differences and more closely reflect complexities seen in vivo. Another key distinction between the ABM and the ODE model is that the ABM is discrete and stochastic, whereas the ODE model is continuous and deterministic. These properties of the ABM might have caused it to wander away from locally stable equilibrium, leading to what we observed in Figure 3C and D. Tumor growth reach equilibrium faster in the ABM than in the ODE model, leading to the lack of intermediate-sized tumors on Day 19 in the ABM. Nevertheless, the enhanced granularity and versatility of ABMs come at the cost of longer computational time and increased difficulty in parameterizing and analyzing the model. Because the ABM updates each cell individually at each time step, simulations slow down significantly when the number of tumor cells increases exponentially. Thus, simulating tumor and immune dynamics at a realistic scale is computationally prohibitive. ABMs generally have many more parameters than the ODE model, making parametrization of the model challenging. In the ODE, we used sensitivity analysis to determine which parameters impact the tumor outcome most and focused our calibration and analytical efforts on those. Sensitivity analysis of ABM parameters, though possible [3, 34], is no trivial task. Future avenues of exploration include using machine learning to overcome the shortcomings of ABMs. In our upcoming work, we plan to combine this ABM with machine learning algorithms to predict the tumor-immune landscape after immune checkpoint blockade, which can make simulating larger virtual cohorts or larger number of cells more feasible.

## 5 Conclusions

We presented the first side-by-side comparison of an ABM with an ODE model for ICIs targeting the PD1/PD-L1 immune checkpoint. We simulated the responses to immune checkpoint blockade therapy in a virtual cohort with diverse tumor-immune characteristics. In particular, we emphasized the importance of including spatial components in mathematical models of immunotherapy by elucidating the additional insights that the ABM provided regarding the spatial complexity of the TME and their impact on therapeutic outcomes. Our computational method can efficiently enhance discovery of key spatial elements, inform biomarker development and validate findings from ongoing clinical data. Even though our model was built for ICIs and was calibrated with in vivo bladder cancer data, our modeling framework and methodology can be applied to other cancers or other forms of cancer immunotherapy.

