## Supplemental Materials for "Agent-Based Modeling of Virtual Tumors Reveals the Critical Influence of Microenvironmental Complexity on Immunotherapy Efficacy"

Jun 28, 2024

#### 1 Supplemental Figures

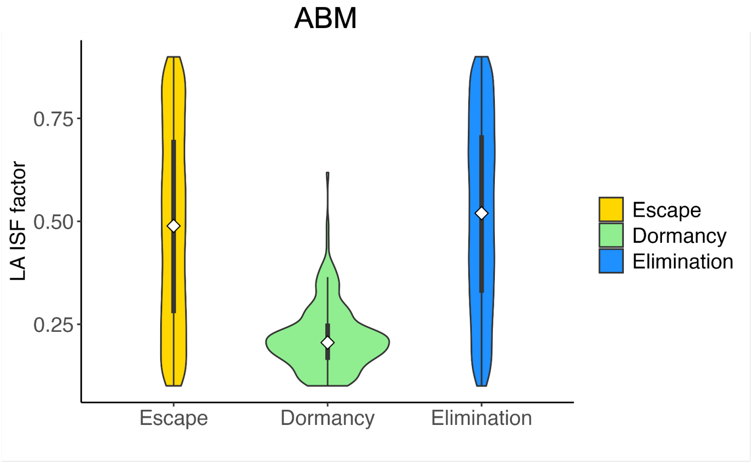


Figure 1: Violin plot of the distribution of LA ISF factor ($r_{\text{ISF}}$) in the virtual cohort associated with each outcome, with the shape showing probability density, the white circle showing the median, and the black lines showing the interquartile range. Elimination (blue): tumor size $< 0.1\text{mm}^3$; Dormancy (green): $0.1 <$ tumor size $< 500\text{mm}^3$; Escape (yellow): tumor size $> 500\text{mm}^3$.

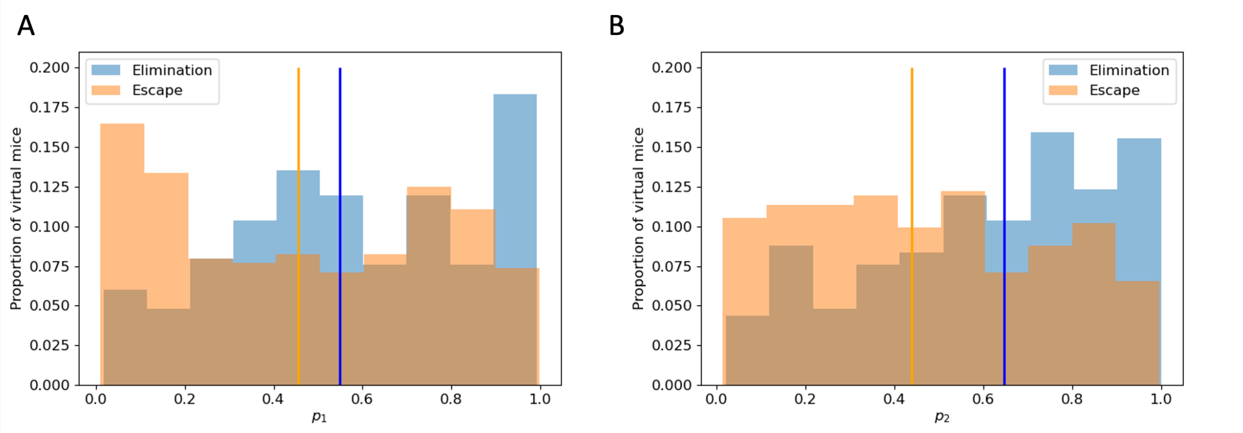


Figure 2: Histogram of $p_1$ and $p_2$ in"elimination" and "escape" groups respectively, with the y-value normalized by the total number of virtual mice in that group. Vertical lines: the median $\mu$ of virtual mice in each group. $p_1$: probability of HA tumor cell death via fast killing; $p_2$: probability of LA tumor cell death via fast killing.

### 2 Supplemental Tables

| Name | Description | Value(s) | Source/Notes |
|---|---|---|---|
| | Tumor Cell Parameters | | |
| $\alpha_n$ | proliferation rate | $2.6\,\mathrm{d}^{-1}$ | Calibrated |
| $O_T^{\text{prolif}}$ | Maximum number of occupied neighbors that still allows tumor cell proliferation (out of 26) | 20 | Calibrated |
| $\delta_n$ | Apoptosis rate | $0.05\,\mathrm{d}^{-1}$ | Estimated [1] |
| $K_{\text{max}}$ | Maximum total number of tumor cells allowed | 60000 | Assumed |
| $K_{\text{edge}}$ | Maximum number of tumor cells allowed to touch the boundary | 36 | Assumed |
| | Immune Cell Parameters | | |
| $\mu$ | Tumor-induced recruitment to TME | $8\ \mathrm{d}^{-1}$ | Calibrated |
| $\alpha_t$ | Base proliferation rate | $0\,\mathrm{d}^{-1}$ | Assumed |
| $\alpha_{nt}$ | Max ISF-stimulated CTL proliferation rate | $0.15\,\mathrm{d}^{-1}$ | [5, 10, 9] |
| $\delta_t$ | Apoptosis rate | $0.05\,\mathrm{d}^{-1}$ | [5, 9] |
| $\beta$ | Immune cell conjugation rate | $1.2\,\mathrm{h}^{-1}$ | [7, 1] |
| $m$ | Movement rate | $2\,\mu\mathrm{m}\,\mathrm{min}^{-1}$ | Estimated [1] |

| | | | |
|---|---|---|---|
| $n_{\text{move}}$ | Number of consecutive movement steps attempted when an immune cell moves | 4 | Estimated [1] |
| $\delta_{\text{exhaust}}$ | CTL exhaustion rate | 0.01 $d^{-1}$ | Estimated |
| | Immune Stimulatory Factor Parameters | | |
| $r_{\text{ISF}}$ | ISF expression by LA tumor cells compared to HA tumor cells | 0.5 | Assumed [1] |
| $\gamma_I$ | EC50 for ISF stimulation of CTL proliferation | 10 | Assumed |
| $n_I$ | Hill coefficient for ISF stimulation of CTL proliferation | 2 | Estimated [1] |
| $f_I$ | Factor determining maximal possible increase to immune proliferation due to ISF | 2.5 | Estimated [1] |
| $\gamma_m$ | EC50 for magnitude of ISF gradient affecting immune cell movement along gradient | 2 | Estimated [1] |
| $n_m$ | Hill coefficient for magnitude of ISF gradient affecting immune cell movement along gradient | 2 | Estimated [1] |
| $a_{\text{reach}}$ | Maximum reach of ISF from one tumor cell in any one direction | 5 | Estimated [1] |
| | Cell-kill Parameters | | |
| $\delta_{\text{slow}}$ | Slow-killing rate | $12\,d^{-1}$ | [4] |
| $\delta_{\text{fast}}$ | Fast-killing rate | $48\,d^{-1}$ | [4] |
| $p_1$ | Probability of HA tumor cell death via fast killing | 0.92 | Assumed [11] |
| $p_2$ | Probability of HA tumor cell death via fast killing | 0.33 | Assumed [11] |
| | PD-1/PD-L1 Parameters | | |
| $\rho_P$ | Concentration of PD-1 on CTLs | 0.6426 nM | [3] |

| | | | |
|---|---|---|---|
| $k_{f_5}$ | Association rate of PD-1-PD-L1 reaction | $100\,\mathrm{nM}^{-1}\,\mathrm{d}^{-1}$ | [6] |
| $k_{r_5}$ | Dissociation rate of PD-1-PD-L1 reaction | $8.25 \times 10^{5}\,\mathrm{d}^{-1}$ | [6] |
| $\gamma_e$ | EC50 of PD-1-PD-L1 complex effects on immune cells | $7.53 \times 10^{-4}\,\mathrm{nM}$ | Computed [1] |
| Miscellaneous Parameters | | | |
| $\Delta t$ | Tumor update duration | $15\,\mathrm{min}$ | Chosen |
| $\Delta t_{\mathrm{imm}}$ | Immune update duration | $7.5\,\mathrm{min}$ | Chosen |
| $L_{G0}$ | Length of time in $G_0$ during which cells cannot proliferate | $9\,\mathrm{h}$ | [8] |
| $h$ | Distance between adjacent voxels | $20\,\mu\mathrm{m}$ | One cell width [1] |
| $O_I^{\mathrm{prolif}}$ | Maximum number of occupied neighbors that still allows immune cell proliferation (out of 26) | 22 | Assumption [1] |
| $O_{\mathrm{max}}^{\mathrm{move}}$ | Maximum number of occupied neighbors that still allows movement (out of 26) | 25 | Assumption [1] |

Table 1: All ABM parameters.

| Parameter | Description | Value (Baseline) | Units | Source |
|---|---|---|---|---|
| $\alpha_n$ | Proliferation rate of high antigen tumor cells | 0.498 | per day | Calibrated |
| $\alpha_m$ | Proliferation rate of low antigen tumor cells | 0.498 | per day | Calibrated |
| K | Carrying capacity for tumor cells | $2394 \times 10^6$ | # of cells | Calibrated |
| $\delta_{ns}$ | Maximum CTL-induced death rate of high antigen tumor cells via the slow killing mechanism | 1 - 12 (4) | per day | Estimated ([2]) |
| $\delta_{ms}$ | Maximum CTL-induced death rate of low antigen tumor cells via the slow killing mechanism | 1 - 12 (4) | per day | Estimated ([2]) |
| $\delta_{nf}$ | CTL-induced death rate of high antigen tumor cells via the fast-killing mechanism | 1e-8 to 1e-6 ($2.5 \times 10^{-7}$) | per cell per day | Estimated ([10]) |
| $\delta_{mf}$ | CTL-induced death rate of low antigen tumor cells via the fast-killing mechanism | 1e-8 to 1e-6 ($2.5 \times 10^{-7}$) | per cell per day | Estimated ([10]) |
| $\mu$ | Activation and recruitment rate of T cells | 5e3 to 1.5e5 ($2 \times 10^4$) | # per day | Estimated |
| $p_1$ | Probability of high antigen tumor cells death via the fast-killing mechanism | 0 to 1 (0.92) | dimensionless | Estimated |
| $p_2$ | Probability of low antigen tumor cells death via the fast-killing mechanism | 0 to 1 (0.33) | dimensionless | Estimated |
| $\alpha_{nt}$ | Maximum rate of CTL proliferation activated by N cells | 0 to 0.5 (0.15) | per day | Estimated |
| $\alpha_{mt}$ | Maximum rate of CTL proliferation activated by M cells | 0 to 0.5 (0.15) | per day | Estimated |

Table 2: Selective ODE parameters

[3] Xiaoxiao Cheng, Vaclav Veverka, Anand Radhakrishnan, Lorna C Waters, Frederick W Muskett, Sara H Morgan, Jiandong Huo, Chao Yu, Edward J Evans, Alasdair J Leslie, et al.

Structure and interactions of the human programmed cell death 1 receptor. *Journal of Biological Chemistry*, 288(17):11771–11785, 2013.
